# Is myeloid-derived growth factor a ligand of the sphingosine-1-phosphate receptor 2?

**DOI:** 10.1101/2024.02.23.581834

**Authors:** Yong-Shan Zheng, Ya-Li Liu, Zeng-Guang Xu, Cheng He, Zhan-Yun Guo

## Abstract

Secretory myeloid-derived growth factor (MYDGF) exerts beneficial effects on organ repair, probably via a plasma membrane receptor; however, the identity of the expected receptor has remained elusive. In a recent study, MYDGF was reported as an agonist of the sphingosine-1-phosphate receptor 2 (S1PR2), an A-class G protein-coupled receptor that mediates the functions of the signaling lipid, sphingosine-1-phosphate (S1P). In the present study, we conducted living cell-based functional assays to test whether S1PR2 is a receptor for MYDGF. In the NanoLuc Binary Technology (NanoBiT)-based β-arrestin recruitment assay and the cAMP-response element (CRE)-controlled NanoLuc reporter assay, S1P could efficiently activate human S1PR2 overexpressed in human embryonic kidney (HEK) 293T cells; however, recombinant human MYDGF, overexpressed either from *Escherichia coli* or HEK293 cells, had no detectable effect. Thus, the results demonstrated that human MYDGF is not a ligand of human S1PR2. Considering the high conservation of MYDGF and S1PR2 in evolution, MYDGF is also probably not a ligand of S1PR2 in other vertebrates.

## 1. Introduction

Myeloid-derived growth factor (MYDGF), formerly known as chromosome 19 open reading frame 10 (C19orf10), is widely present across the animal kingdom and is relatively conserved in evolution. In 2015, Korf-Klingebiel et al. first reported that MYDGF can promote cardiac myocyte survival and angiogenesis after myocardial infarction in mice [1]. Since then, more functions of MYDGF have been reported in various animal models, such as protection against pressure overload-induced heart failure and improvement of heart regeneration [2,3]; protection against kidney injury in ischemia/reperfusion and some kidney diseases [4–7]; alleviation of endothelial injury and protection against atherosclerosis [8,9]; alleviation of non-alcoholic fatty liver disease [10]; maintenance of glucose homeostasis in diabetes [11]; and regulation of neutrophil motility and suppression of vascular smooth muscle cell dedifferentiation [12,13]. Thus, it seems that MYGDF has beneficial effects on various organs and tissues, especially under stressed conditions.

According to the solved solution or crystal structure [14,15], mature human MYDGF (142 amino acids) folds into a unique β-sandwich structure composed of 10 β-strands. Despite its name, MYDGF is widely expressed in various human cells, according to the recently published single cell RNA sequencing data (www.proteinatlas.org/ENSG00000074842-MYDGF/single+cell+type). *In vivo*, MYDGF is synthesized as a precursor containing an N-terminal signal peptide and a C-terminal endoplasmic reticulum (ER)-retention motif. As a result, most of the mature MYDGF is retained in ER and Golgi apparatus, with unknown functions [16]. Only a small fraction of mature MYDGF is secreted and functions as an extracellular factor. The median plasma MYDGF concentration is approximately 3.3 ng/ml (or 0.2 nM) in healthy human adults, and is increased by approximately 2.7-fold in patients with acute myocardial infarction [17]. Administration of recombinant MYDGF had beneficial effects in animal models and cultured cells [1–5,7–13], implying that extracellular MYDGF has biological functions.

It is hypothesized that extracellular MYDGF exerts its function via a plasma membrane receptor. However, the identity of the expected receptor is unknown. A recent study reported that sphingosine-1-phosphate receptor 2 (S1PR2) is the long-sought MYDGF receptor [13]. S1PR2 is an A-class G protein-coupled receptor (GPCR) that uses sphingosine-1-phosphate (S1P) as its endogenous agonist. The signaling lipid S1P and its five cognate receptors (S1PR1–5) have important functions in various physiological and pathological conditions [18–22]. In the present study, we tested whether MYDGF is a ligand of S1PR2 using two living cell-based functional assays. In the NanoLuc Binary Technology (NanoBiT)-based β-arrestin recruitment assay and the cAMP-response element (CRE)-controlled NanoLuc reporter assay, the signaling lipid S1P could efficiently activate human S1PR2. However, recombinant human MYDGF, produced either from *Escherichia coli* or human embryonic kidney (HEK) 293 cells, had no detectable effect, suggesting that MYDGF is not a ligand of S1PR2. In the future, more studies are needed to identify the real receptor for MYDGF.

## 2. Materials and methods

### 2.1. Bacterial overexpression of human MYDGF

The DNA sequence encoding a designed human MYDGF precursor was chemically synthesized at Tsingke Biotechnology (Shanghai, China) using *E. coli*-biased codons. After cleavage with restriction enzyme NdeI and EcoRI, the synthetic DNA fragment was ligated into a pET vector, resulting in the expression construct pET/6×His-EK-MYDGF that encodes a precursor carrying a 6×His-tag and an enterokinase cleavage site at the N-terminus of the mature human MYDGF (Fig. S1). Its nucleotide sequence was confirmed by DNA sequencing at Tsingke Biotechnology (Shanghai, China).

Thereafter, the expression construct was transformed into the *E. coli* strain Shuffle T7, and the bacteria were cultured in liquid Luria-Bertani medium (plus 100 μg/ml ampicillin) to OD_600_ ≈ 1.0 at 37 °C with vigorous shaking. Subsequently, isopropyl β-D-thiogalactoside (IPTG) was added to the final concentration of 0.5 mM, and the bacteria were continuously cultured at 20 °C overnight. After harvested by centrifugation (5000 *g*, 10 min), the bacteria pellet was suspended in lysis buffer (50 mM Tris-Cl, 100 mM NaCl, pH 7.4) and lysed by sonication. After centrifugation (15000 *g*, 30 min), the supernatant was applied to an immobilized metal ion affinity chromatography (Ni^2+^ column), and the fraction of 6×His-EK-MYDGF was eluted by 250 mM imidazole. After dialysis against 20 mM Tris-Cl buffer (pH 8.0), the MYDGF precursor fraction was subjected to an ion-exchange chromatography. After eluted from the DEAE column by a NaCl gradient, the purified MYDGF precursor was treated with recombinant enterokinase (Yaxin Bio, Shanghai, China) at 25 °C for 16 h to remove the N-terminal tag. The recombinant MYDGF samples at each preparation step were analyzed by sodium dodecyl sulfate-polyacrylamide gel electrophoresis (SDS-PAGE). Molecular mass of the mature MYDGF was measured on an AB SCIEX TripleTOF 5600 mass spectrometer at BiotechPack Scientific (Beijing, China).

### 2.2. Generation of the expression constructs for human S1PR2

The human genomic DNA was extracted from HEK293T cells and used as the template for amplification of the coding region of human S1PR2 via polymerase chain reaction. After amplification using pfu DNA polymerase and appropriate oligo primers (Table S1), the DNA fragment was cleaved by restriction enzyme AgeI, and ligated into cleaved vectors containing a blunt end and a sticky end. As shown in Fig. S2, the resultant construct pcDNA6/S1PR2 encodes an untagged human S1PR2, and the construct pTRE3G-BI/S1PR2-LgBiT:SmBiT-ARRB2 co-expresses a C-terminally large NanoLuc fragment (LgBiT)-fused human S1PR2 and an N-terminally small complementation tag (SmBiT)-fused human β-arrestin 2 (ARRB2). Their nucleotide sequence was confirmed by DNA sequencing at Tsingke Biotechnology (Shanghai, China).

### 2.3. The NanoBiT-based β-arrestin recruitment assay

HEK293T cells were transiently cotransfected with the expression construct pTRE3G-BI/S1PR2-LgBiT:SmBiT-ARRB2 and the expression control vector pCMV-TRE3G (Clontech, Mountain View, CA, USA) using the transfection reagent Lipo8000 (Beyotime Technology, Shanghai, China). Next day, the transfected cells were trypsinized, suspended in the complete DMEM medium with 4 ng/ml of doxycycline (Dox), and seeded into a white opaque 96-well plate. After cultured at 37 °C for ∼24 h to ∼90% confluence, the medium was removed and the pre-warmed activation solution (serum-free DMEM plus 1% BSA) containing NanoLuc substrate was added to the living cells (40 μl/well, containing 1.0 μl of NanoLuc substrate stock from Promega, Madison, WI, USA). Thereafter, bioluminescence data were collected for ∼5 min on a SpectraMax iD3 plate reader (Molecular Devices, Sunnyvale, CA, USA). Subsequently, solution of S1P (MedChemExpress, Shanghai, China) or recombinant MYDGF was added to these wells (10 μl/well, diluted in the activation solution), and bioluminescence data were continuously collected for ∼10 min. The recombinant MYDGF was produced either from *E. coli* in our present study or HEK293 cells in Sino Biological (Beijing, China). The *E. coli*-derived human MYDGF is a mature form and the HEK293-secreted human MYDGF contains a 6×His-tag at its C-terminus. If necessary, the measured absolute bioluminescence signals were corrected for inter well variability by forcing all curves after addition of NanoLuc substrate (without ligand) to same level. For the S1P dose curve, the measured bioluminescence values at 649 s were plotted with S1P concentrations using SigmaPlot 10.0 software (SYSTAT software, Chicago, IL, USA).

### 2.4. The CRE-controlled NanoLuc reporter assay

HEK293T cells were transiently cotransfected with the expression construct pcDNA6/S1PR2 and a NanoLuc reporter vector pNL1.2/CRE using the transfection reagent Lipo8000 (Beyotime Technology). Next day, the transfected cells were trypsinized and seeded into a white opaque 96-well plate. After cultured in complete DMEM medium for ∼24 h to ∼90% confluence, the medium was removed and the activation solution (serum-free DMEM plus 1% BSA) containing varied concentrations of S1P (MedChemExpress) or recombinant MYDGF produced either from *E. coli* (our present work) or HEK293 cells (from Sino Biological) was added (50 μl/well). After continuously cultured at 37 °C for 4 h, bioluminescence was measured on a SpectraMax iD3 plate reader (Molecular Devices) after addition of the diluted NanoLuc substrate (10 μl/well, 30-fold dilution of the Promega substrate stock using the activation solution). The measured bioluminescence data were expressed as mean ± standard deviate (SD, *n* = 3) and plotted using SigmaPlot10.0 software (SYSTAT software).

## 3. Results

### 3.1. Preparation of recombinant human MYDGF via bacterial overexpression

Mature human MYDGF (142 residues) forms a β-sandwich structure with 10 β-strands [14,15]. Among them, the third and fifth β-strands are tethered together by a highly conserved intramolecular disulfide bond. To produce the correctly folded MYDGF protein, we employed the *E. coli* strain Shuffle T7 because it can promote disulfide bond formation in the cytosol. To facilitate purification, a 6×His-tag was introduced into the N-terminus of mature human MYDGF (Fig. S1). To remove the N-terminal tag after purification, an enterokinase cleavage site was introduced between the tag and the mature MYDGF (Fig. S1).

After the transformed Shuffle T7 bacteria were induced by IPTG, an additional protein band (indicated by an asterisk), with an apparent molecular weight of slightly less than 20 kDa, appeared on SDS-PAGE (Fig. 1A), consistent with the theoretical value (17.5 kDa) of the overexpressed 6×His-EK-MYDGF. After sonication, most of the overexpressed MYDGF precursor was present in the supernatant and was conveniently purified using an Ni^2+^ column, as analyzed using SDS-PAGE (Fig. 2A). After dialysis, the MYDGF precursor was further purified using a DEAE ion-exchange column and then treated with enterokinase to remove the N-terminal tag (Fig. S1). After enterokinase treatment, a slightly smaller band (indicated by an octophone) appeared on SDS-PAGE (Fig. 1B), suggesting removal of the N-terminal tag. The recombinant mature human MYDGF displayed a molecular mass of 15833.6 as measured by mass spectrometry (Fig. S3), consistent with its theoretical value (15832.6). Thus, mature human MYDGF could be conveniently prepared via bacteria overexpression and *in vitro* maturation.

**Fig. 1.**
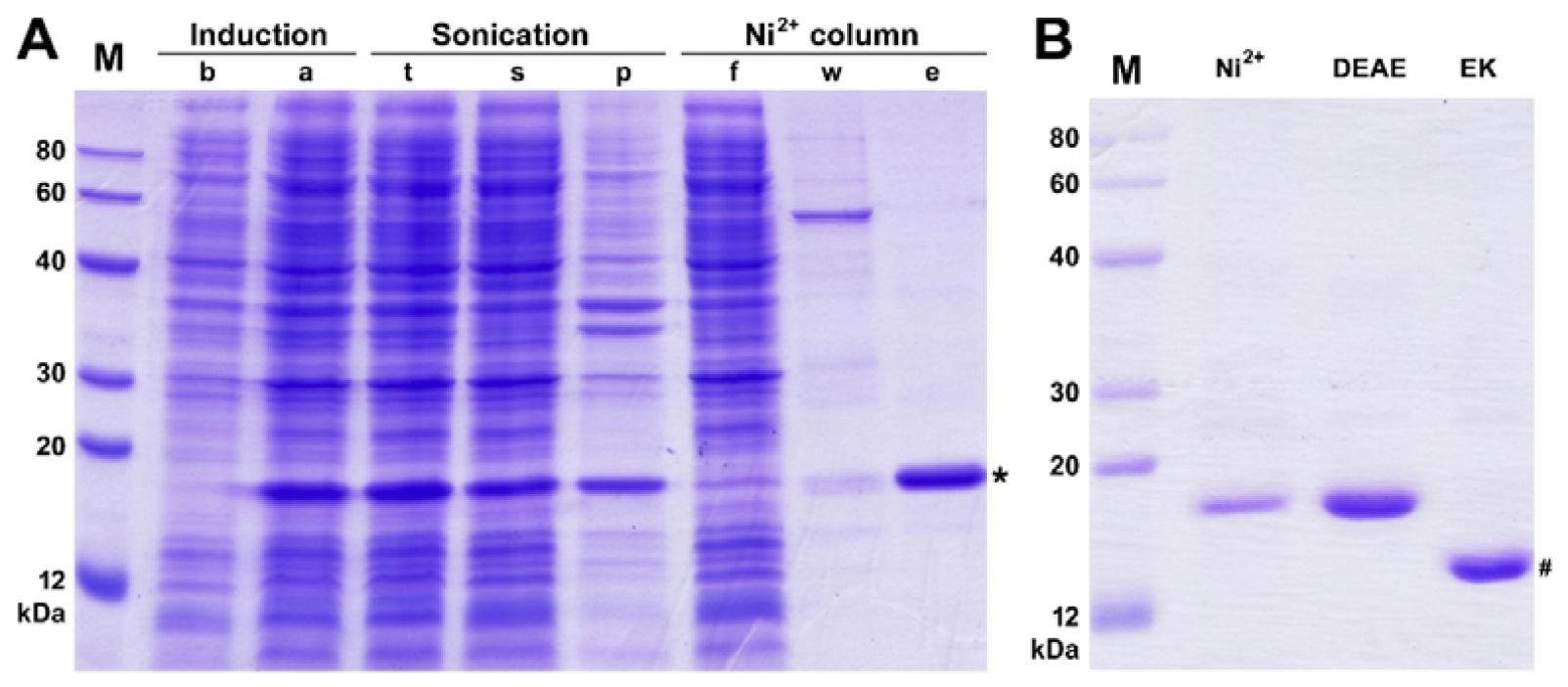
Preparation of mature human MYDGF via bacteria overexpression. (A) SDS-PAGE analysis of the overexpressed MYDGF precursor at different preparation steps. Lane b, before IPTG induction; lane a, after IPTG induction; lane t, total lysate after sonication; lane s, supernatant of the lysate; lane p, pellet of the lysate; lane f, low-through from the Ni^2+^ column; lane w, washing fraction by 50 mM imidazole; lane e, eluted fraction by 250 mM imidazole. Band of the MYDGF precursor is indicated by an asterisk. (B) SDS-PAGE analysis of the overexpressed precursor and mature MYDGF. Lane Ni^2+^, the precursor eluted from the Ni^2+^ column; lane DEAE, the precursor eluted from the DEAE column; lane EK, the precursor treated with enterokinase. Band of the mature MYDGF is indicated by an octophone.

**Fig. 2.**
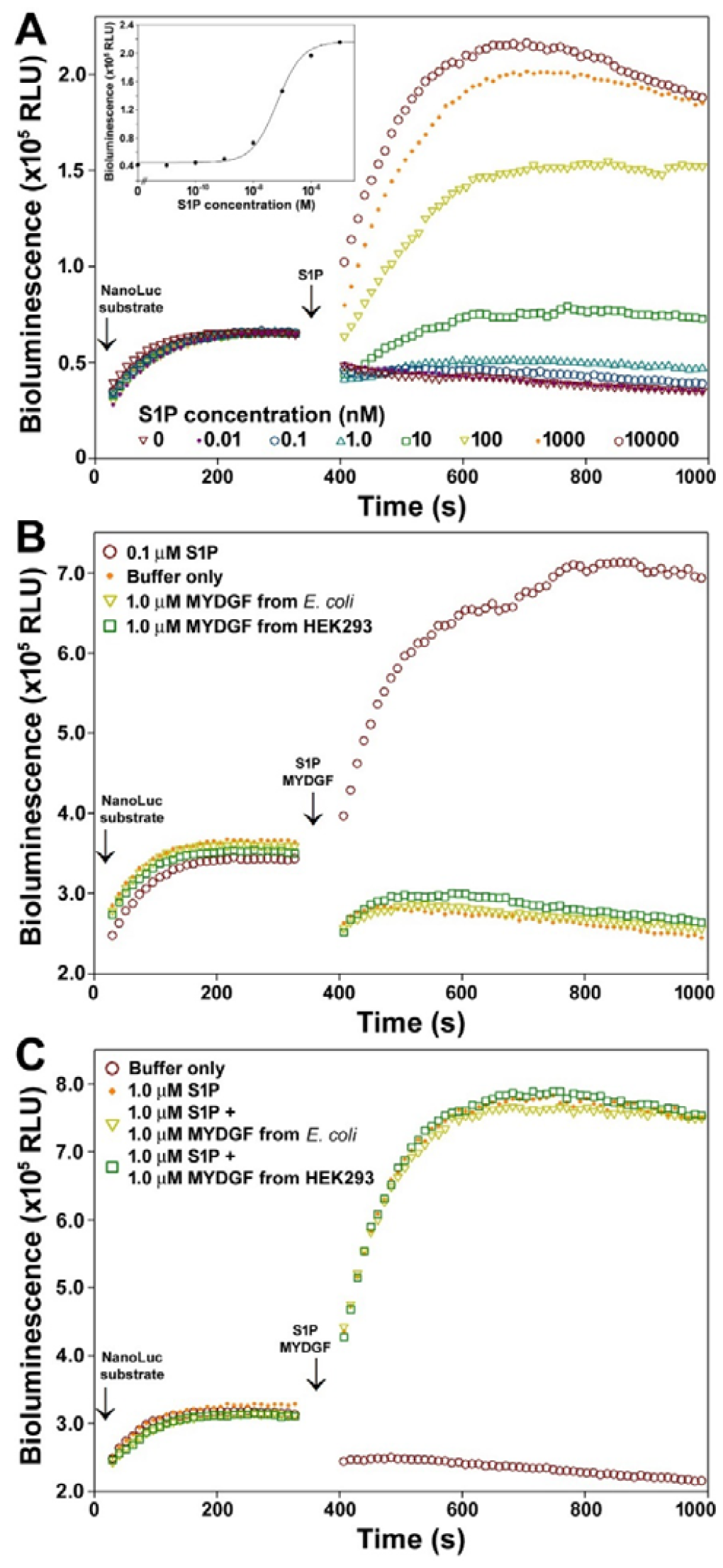
The NanoBiT-based β-arrestin recruitment assay for human S1PR2. (**A**) Measured bioluminescence data after sequential addition of the NanoLuc substrate and different concentrations of S1P to living HEK293T cells coexpressing S1PR2-LgBiT and SmBiT-ARRB2. Inner panel, dose curve of S1P activating human S1PR2 measured by the β-arrestin recruitment assay. The measured bioluminescence data at 649 s in panel A were plotted with S1P concentrations using SigmaPlot 10.0 software. (**B**) Effect of the recombinant human MYDGF produced either from *E. coli* or HEK293 cells on β-arrestin recruitment to human S1PR2. (**C**) Effect of the recombinant human MYDGF produced either from *E. coli* or HEK293 cells on the S1P-induced β-arrestin recruitment to human S1PR2.

We also purchased recombinant human MYDGF from a global leading supplier of bioactive proteins (Sino Biological, Beijing China, Cat: 13469-H08H) for activity assays. Their recombinant human MYDGF was secretorily overexpressed in HEK293 cells and carries a 6×His-tag at the C-terminus.

### 3.2. Effect of MYDGF on S1PR2 activation detected by the β-arrestin recruitment assay

To test whether MYDGF is an agonist of the receptor S1PR2, we first employed the recently developed NanoBiT-based β-arrestin recruitment assay, which has been used to monitor GPCR activation in recent years [23[25]. For this purpose, an inactive LgBiT was genetically fused to the intracellular C-terminus of human S1PR2, and a low-affinity SmBiT complementation tag was genetically fused to the N-terminus of human β-arrestin 2 (ARRB2). If the resultant S1PR2-LgBiT was activated by an agonist, the coexpressed SmBiT-ARRB2 would bind to it, resulting in complementation of the receptor-fused LgBiT with the ARRB2-fused SmBiT via a proximity effect, which would restore luciferase activity.

To coexpress S1PR2-LgBiT and SmBiT-ARRB2 in transfected cells in a controllable manner, we employed the Tet-On inducible expression system with a bi-directional promoter. After transient transfection of HEK293T cells, Dox was used to induce the coexpression of S1PR2-LgBiT and SmBiT-ARRB2. After the addition of the NanoLuc substrate to these induced cells, low bioluminescence was detected (Fig. 2A), implying background complementation of the ARRB2-fused SmBiT with the S1PR2-fused LgBiT. After the further addition of the endogenous agonist S1P, the measured bioluminescence increased quickly in a dose-dependent manner (Fig. 2A), implying that S1PR2-LgBiT was activated by S1P and bound to SmBiT-ARRB2. According to the maximal bioluminescence induced by different concentrations of S1P, a typical sigmoidal receptor activation curve was obtained (Fig. 2A, inner panel), with a calculated EC_50_ value of approximately 68 nM. Thus, the NanoBiT-based β-arrestin recruitment assay could sensitively monitor S1PR2 activation in the transfected HEK293T cells.

In contrast, the addition of recombinant MYDGF at concentrations as high as 1.0 μM (either produced from *E. coli* or HEK293 cells) had no significant activation effect in the β-arrestin recruitment assay (Fig. 2B), implying that human MYDGF is not an agonist for human S1PR2. The addition of recombinant MYDGF also had no significant effect on S1PR2 activation induced by S1P (Fig. 2C), suggesting that human MYDGF is also not an antagonist for human S1PR2.

### 3.3. Effect of MYDGF on S1PR2 activation detected by the CRE-controlled NanoLuc reporter assay

We further monitored the activation of S1PR2 using a CRE-controlled NanoLuc reporter assay, because this receptor can couple to the Gs signaling pathway and increase the intracellular cAMP level after activation [26]. To conduct this assay, the expression construct pcDNA6/S1PR2, encoding an untagged human S1PR2, and the reporter vector pNL1.2/CRE, were transiently cotransfected into HEK293T cells. After these cells were treated with S1P, the measured bioluminescence increased in a typical sigmoidal manner (Fig. 3A), with a calculated EC_50_ value of approximately 71 nM, implying that S1P can efficiently activate S1PR2. S1P had no significant effect on HEK293T cells transfected only with the NanoLuc reporter vector (Fig. 3A, inner panel), implying that HEK293T cells did not express endogenous S1P receptors. Thus, the CRE-controlled NanoLuc reporter assay could be used to monitor the activation of S1PR2 in the transfected HEK293T cells.

**Fig. 3.**
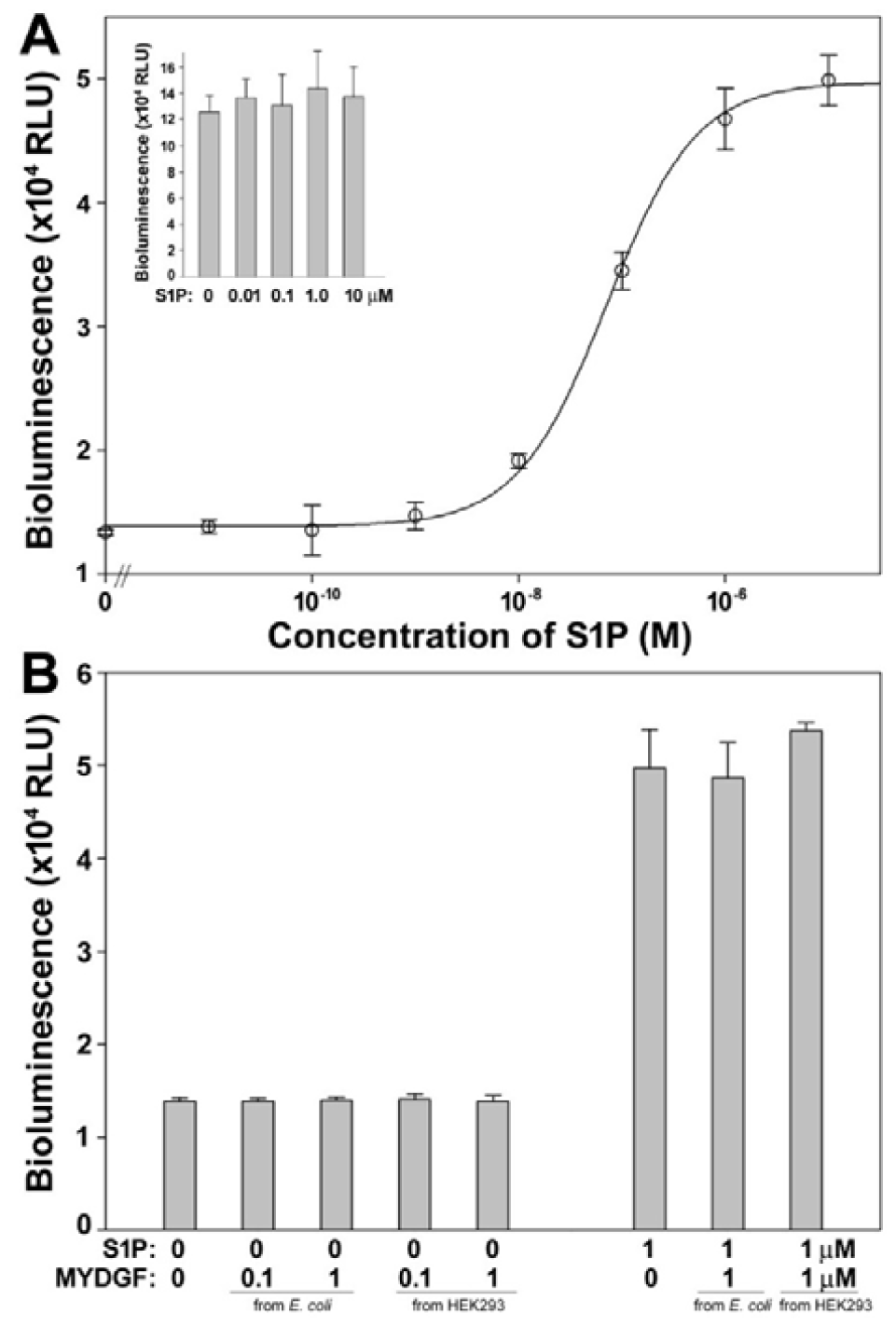
The CRE-controlled NanoLuc reporter assay for human S1PR2. (A) Activation of human S1PR2 by S1P monitored by the CRE-controlled NanoLuc reporter. Inner panel, effect of S1P on HEK293T cells transfected with the reporter vector pNL1.2/CRE only. (**B**) Effect of the recombinant human MYDGF on human S1PR2 activation under absence or presence of S1P.

The addition of recombinant MYDGF (produced either from *E. coli* or HEK293 cells) had no detectable activation effect in the CRE-controlled NanoLuc reporter assay (Fig. 3B), suggesting that human MYDGF has no agonistic function towards human S1PR2. Recombinant MYDGF also did not affect S1PR2 activation induced by S1P (Fig. 3B), suggesting that human MYDGF has no antagonistic function on human S1PR2. Thus, the recombinant human MYDGF had not detectable agonistic or antagonistic effect on human S1PR2 in both the NanoBiT-based β-arrestin recruitment assay and the CRE-controlled NanoLuc reporter assay. This suggested that human MYDGF is not a ligand of human S1PR2.

## 4. Discussion

In the present study, we tested whether S1PR2 is the receptor of MYDGF using living cell-based functional assays. In the NanoBiT-based β-arrestin recruitment assay and the CRE-controlled NanoLuc reporter assay, we could not detect any agonistic or antagonistic effect of recombinant human MYDGF on human S1PR2, although the lipid agonist S1P efficiently activated this receptor in both assays. Thus, our present results demonstrated that human MYDGF is not a ligand of human S1PR2. As shown in Fig. S4C,D, both MYDGF and S1PR2 display high amino acid sequence homology among vertebrates, suggesting that their functions are highly conserved in evolution. Thus, MYDGF is unlikely to be a ligand of S1PR2 in other vertebrates.

In the recent paper [13], the authors first speculated that S1PR2 might be the receptor of MYDGF, based only on the previous observation that the S1PR2 signaling pathway is implicated in dedifferentiation, proliferation, and migration of vascular smooth muscle cells. Thereafter, the authors conducted molecular docking to test whether MYDGF binds to S1PR2; however, they never mentioned which species their S1PR2 and MYDGF proteins came from [13]. We thought that their proteins were most likely from human (*Homo sapiens*), mouse (*Mus musculus*), or rat (*Rattus norvegicus*) (Fig. S4A,B). In their docking analysis, they identified three strange interaction pairs between MYDGF and S1PR2: the smallest Gly75 of MYDGF versus the small Ala263 of S1PR2, the positively charged His82 of MYDGF versus the positively charged Lys269 of S1PR2, and the negatively charged Glu122 of MYDGF versus the hydrophobic Leu176 of S1PR2 [13]. Considering the above-mentioned residues, mouse or rat MYDGF and S1PR2 were likely used in their molecular docking analysis (Fig. S4A,B). For the ligand MYDGF, the three so-called S1PR2-binding residues are not conserved among different vertebrates, or even among mammals (Fig. S4C). For the receptor S1PR2, the three so-called MYDGF-binding residues are also variable among different vertebrates (Fig. S4D). In general, residues for ligand-receptor interactions are typically highly conserved in both the ligand and the receptor. Thus, it seems that their docking analysis was not reliable.

After molecular docking, the authors analyzed the interaction between purified MYDGF and S1PR2 via surface plasmon resonance (SPR) [13]. However, they never mentioned where their S1PR2 sample came from or how it was prepared [13]. All GPCRs, including S1PR2, as integral membrane proteins with seven transmembrane domains, are extremely difficult to prepare. Once solubilized from the plasma membrane using detergents, they are prone to inactivation in subsequent purification, storage, or immobilization steps. The authors never tested whether their purified S1PR2 was active, such as by assessing binding to its endogenous agonist, S1P [13]. Moreover, the transmembrane domain and the intracellular domain are exposed after solubilization from the plasma membrane. Thus, the so-called interaction between MYDGF and S1PR2 detected by SPR or by co-immunoprecipitation (co-IP) have alternative explanations: it might have been caused by nonspecific interaction of MYDGF with a denatured receptor or with a receptor patch that is only accessible in the detergent-solubilized receptor.

To test whether MYDGF is an agonist of S1PR2, the most convincing experiments are living cell-based functional assays using the endogenous agonist S1P as a positive control. Unfortunately, the authors did not conduct these essential assays to support their claim [13]. In recent years, research into the deorphanization of orphan receptors or orphan ligands has slowed down, and some so-called deorphanization reports cannot be reproduced by other laboratories, such as GPR83 versus the peptide PEN and proCCK56-63 [27[31], GPR146 versus the proinsulin C-peptide [32,33], GPR160 versus the neuropeptide CART [34,35], and GPR35 versus CXCL17 [36[38]. Successful deorphanization depends on rationally designed experiments and strong evidence, otherwise the claim might represent hype rather than hope for the scientific community.

## Author contributions

YSZ conducted the experiments. YLL and ZGX analyzed the data. CH and ZYG directed the research and wrote the paper.

## Data availability statement

The data supporting this study are available in the supplementary materials of this article.

## Acknowledgements

This work was supported by grant from the National Natural Science Foundation of China (grant no. 31971193).

