## Supplemental Table and Figures for "Is myeloid-derived growth factor a ligand of the sphingosine-1-phosphate receptor 2?"

**Table S1.** Synthetic oligo primers for amplification of human S1PR2.

| Oligo name | Nucleotide sequence (5' to 3') | Restriction site | Construct generated |
|---|---|---|---|
| S1PR2-F-Blunt | ATG GGC AGC TTG TAC TCG GAG TAC CT | no | pcDNA6/S1PR2 |
| S1PR2-R-AgeI | A ACC GGT GG GAC CAC CGT GTT GCC CTC CAG AAA | AgeI | |
| S1PR2-F-Blunt | ATG GGC AGC TTG TAC TCG GAG TAC CT | no | pTRE3G-BI/<br>S1PR2-LgBiT:SmBiT-ARRB[illegible] |
| S1PR2-LgBiT-AgeI | GA ACC GGT GAC CAC CGT GTT GCC CTC CAG | AgeI | |

```
                                   NdeI
  1   ATG CAT CAC CAT CAC CAC CAT ATG GAT GAC GAT GAC AAA GTG TCC GAG CCG ACG ACG GTG GCG TTT GAC GTG CGT
      TAC GTA GTG GTA GTG GTG GTA TAC CTA CTG CTA CTG TTT CAC AGG CTC GGC TGC TGC CAC CGC AAA CTG CAC GCA
       M   H   H   H   H   H   H   M   D   D   D   D   K   V   S   E   P   T   T   V   A   F   D   V   R

 76   CCA GGC GGC GTC GTG CAT TCT TTC TCA CAT AAC GTG GGC CCG GGT GAC AAA TAT ACG TGT ATG TTC ACT TAC GCC
      GGT CCG CCG CAG CAC GTA AGA AAG AGT GTA TTG CAC CCG GGC CCA CTG TTT ATA TGC ACA TAC AAG TGA ATG CGG
       P   G   G   V   V   H   S   F   S   H   N   V   G   P   G   D   K   Y   T   C   M   F   T   Y   A

151   TCT CAA GGT GGG ACC AAT GAA CAA TGG CAG ATG AGT CTG GGG ACC AGC GAA GAC CAC CAG CAC TTC ACC TGC ACC
      AGA GTT CCA CCC TGG TTA CTT GTT ACC GTC TAC TCA GAC CCC TGG TCG CTT CTG GTG GTC GTG AAG TGG ACG TGG
       S   Q   G   G   T   N   E   Q   W   Q   M   S   L   G   T   S   E   D   H   Q   H   F   T   C   T

226   ATC TGG CGC CCT CAG GGC AAG TCC TAT CTG TAC TTC ACA CAG TTC AAG GCA GAG GTG CGT GGC GCT GAA ATT GAG
      TAG ACC GCG GGA GTC CCG TTC AGG ATA GAC ATG AAG TGT GTC AAG TTC CGT CTC CAC GCA CCG CGA CTT TAA CTC
       I   W   R   P   Q   G   K   S   Y   L   Y   F   T   Q   F   K   A   E   V   R   G   A   E   I   E

301   TAC GCC ATG GCC TAC TCT AAA GCC GCA TTT GAA CGT GAA AGT GAT GTC CCT CTG AAA ACT GAG GAA TTT GAA GTG
      ATG CGG TAC CGG ATG AGA TTT CGG CGT AAA CTT GCA CTT TCA CTA CAG GGA GAC TTT TGA CTC CTT AAA CTT CAC
       Y   A   M   A   Y   S   K   A   A   F   E   R   E   S   D   V   P   L   K   T   E   E   F   E   V

376   ACC AAA ACA GCA GTG GCT CAC CGT CCG GGT GCA TTC AAA GCT GAG CTG TCC AAG CTG GTG ATT GTG GCC AAG GCA
      TGG TTT TGT CGT CAC CGA GTG GCA GGC CCA CGT AAG TTT CGA CTC GAC AGG TTC GAC CAC TAA CAC CGG TTC CGT
       T   K   T   A   V   A   H   R   P   G   A   F   K   A   E   L   S   K   L   V   I   V   A   K   A

                                  EcoRI
451   TCG CGC ACT GAG CTG TAA GAA TTC
      AGC GCG TGA CTC GAC ATT CTT AAG
       S   R   T   E   L   *
```

**Fig. S1.** The nucleotide and amino acid sequences of the MYDGF precursor overexpressed in *E. coli*. The amino acid sequence of mature human MYDGF is shown on red, and that of the enterokinase recognition site is shaded. Enterokinase cleaves the peptide chain at the C-terminus of the DDDDK recognition site. The restriction enzyme cleavage sites for gene cloning are shaded.

### Untagged human S1PR2 in the construct pcDNA6/S1PR2

```
1     ATG GGC AGC TTG TAC TCG GAG TAC CTG AAC CCC AAC AAG GTC CAG GAA CAC TAT AAT TAT ACC AAG GAG ACG CTG
      TAC CCG TCG AAC ATG AGC CTC ATG GAC TTG GGG TTG TTC CAG GTC CTT GTG ATA TTA ATA TGG TTC CTC TGC GAC
       M   G   S   L   Y   S   E   Y   L   N   P   N   K   V   Q   E   H   Y   N   Y   T   K   E   T   L

76    GAA ACG CAG GAG ACG ACC TCC CGC CAG GTG GCC TCG GCC TTC ATC GTC ATC CTC TGT TGC GCC ATT GTG GTG GAA
      CTT TGC GTC CTC TGC TGG AGG GCG GTC CAC CGG AGC CGG AAG TAG CAG TAG GAG ACA ACG CGG TAA CAC CAC CTT
       E   T   Q   E   T   T   S   R   Q   V   A   S   A   F   I   V   I   L   C   C   A   I   V   V   E

151   AAC CTT CTG GTG CTC ATT GCG GTG GCC CGA AAC AGC AAG TTC CAC TCG GCA ATG TAC CTG TTT CTG GGC AAC CTG
      TTG GAA GAC CAC GAG TAA CGC CAC CGG GCT TTG TCG TTC AAG GTG AGC CGT TAC ATG GAC AAA GAC CCG TTG GAC
       N   L   L   V   L   I   A   V   A   R   N   S   K   F   H   S   A   M   Y   L   F   L   G   N   L

226   GCC GCC TCC GAT CTA CTG GCA GGC GTG GCC TTC GTA GCC AAT ACC TTG CTC TCT GGC TCT GTC ACG CTG AGG CTG
      CGG CGG AGG CTA GAT GAC CGT CCG CAC CGG AAG CAT CGG TTA TGG AAC GAG AGA CCG AGA CAG TGC GAC TCC GAC
       A   A   S   D   L   L   A   G   V   A   F   V   A   N   T   L   L   S   G   S   V   T   L   R   L

301   ACG CCT GTG CAG TGG TTT GCC CGG GAG GGC TCT GCC TTC ATC ACG CTC TCG GCC TCT GTC TTC AGC CTC CTG GCC
      TGC GGA CAC GTC ACC AAA CGG GCC CTC CCG AGA CGG AAG TAG TGC GAG AGC CGG AGA CAG AAG TCG GAG GAC CGG
       T   P   V   Q   W   F   A   R   E   G   S   A   F   I   T   L   S   A   S   V   F   S   L   L   A

376   ATC GCC ATT GAG CGC CAC GTG GCC ATT GCC AAG GTC AAG CTG TAT GGC AGC GAC AAG AGC TGC CGC ATG CTT CTG
      TAG CGG TAA CTC GCG GTG CAC CGG TAA CGG TTC CAG TTC GAC ATA CCG TCG CTG TTC TCG ACG GCG TAC GAA GAC
       I   A   I   E   R   H   V   A   I   A   K   V   K   L   Y   G   S   D   K   S   C   R   M   L   L

451   CTC ATC GGG GCC TCG TGG CTC ATC TCG CTG GTC CTC GGT GGC CTG CCC ATC CTT GGC TGG AAC TGC CTG GGC CAC
      GAG TAG CCC CGG AGC ACC GAG TAG AGC GAC CAG GAG CCA CCG GAC GGG TAG GAA CCG ACC TTG ACG GAC CCG GTG
       L   I   G   A   S   W   L   I   S   L   V   L   G   G   L   P   I   L   G   W   N   C   L   G   H

526   CTC GAG GCC TGC TCC ACT GTC CTG CCT CTC TAC GCC AAG CAT TAT GTG CTG TGC GTG GTG ACC ATC TTC TCC ATC
      GAG CTC CGG ACG AGG TGA CAG GAC GGA GAG ATG CGG TTC GTA ATA CAC GAC ACG CAC CAC TGG TAG AAG AGG TAG
       L   E   A   C   S   T   V   L   P   L   Y   A   K   H   Y   V   L   C   V   V   T   I   F   S   I

601   ATC CTG TTG GCC ATC GTG GCC CTG TAC GTG CGC ATC TAC TGC GTG GTC CGC TCA AGC CAC GCT GAC ATG GCC GCC
      TAG GAC AAC CGG TAG CAC CGG GAC ATG CAC GCG TAG ATG ACG CAC CAG GCG AGT TCG GTG CGA CTG TAC CGG CGG
       I   L   L   A   I   V   A   L   Y   V   R   I   Y   C   V   V   R   S   S   H   A   D   M   A   A

676   CCG CAG ACG CTA GCC CTG CTC AAG ACG GTC ACC ATC GTG CTA GGC GTC TTT ATC GTC TGC TGG CTG CCC GCC TTC
      GGC GTC TGC GAT CGG GAC GAG TTC TGC CAG TGG TAG CAC GAT CCG CAG AAA TAG CAG ACG ACC GAC GGG CGG AAG
       P   Q   T   L   A   L   L   K   T   V   T   I   V   L   G   V   F   I   V   C   W   L   P   A   F

751   AGC ATC CTC CTT CTG GAC TAT GCC TGT CCC GTC CAC TCC TGC CCG ATC CTC TAC AAA GCC CAC TAC TTT TTC GCC
      TCG TAG GAG GAA GAC CTG ATA CGG ACA GGG CAG GTG AGG ACG GGC TAG GAG ATG TTT CGG GTG ATG AAA AAG CGG
       S   I   L   L   L   D   Y   A   C   P   V   H   S   C   P   I   L   Y   K   A   H   Y   F   F   A

826   GTC TCC ACC CTG AAT TCC CTG CTC AAC CCC GTC ATC TAC ACG TGG CGC AGC CGG GAC CTG CGG CGG GAG GTG CTT
      CAG AGG TGG GAC TTA AGG GAC GAG TTG GGG CAG TAG ATG TGC ACC GCG TCG GCC CTG GAC GCC GCC CTC CAC GAA
       V   S   T   L   N   S   L   L   N   P   V   I   Y   T   W   R   S   R   D   L   R   R   E   V   L

901   CGG CCG CTG CAG TGC TGG CGG CCG GGG GTG GGG GTG CAA GGA CGG AGG CGG GGC GGG ACC CCG GGC CAC CAC CTC
      GCC GGC GAC GTC ACG ACC GCC GGC CCC CAC CCC CAC GTT CCT GCC TCC GCC CCG CCC TGG GGC CCG GTG GTG GAG
       R   P   L   Q   C   W   R   P   G   V   G   V   Q   G   R   R   R   G   G   T   P   G   H   H   L

976   CTG CCA CTC CGC AGC TCC AGC TCC CTG GAG AGG GGC ATG CAC ATG CCC ACG TCA CCC ACG TTT CTG GAG GGC AAC
      GAC GGT GAG GCG TCG AGG TCG AGG GAC CTC TCC CCG TAC GTG TAC GGG TGC AGT GGG TGC AAA GAC CTC CCG TTG
       L   P   L   R   S   S   S   S   L   E   R   G   M   H   M   P   T   S   P   T   F   L   E   G   N

                        AgeI
1051  ACG GTG GTC CCA CCG GTC TGA TAA
      TGC CAC CAG GGT GGC CAT ACT ATT
       T   V   V   P   P   V   *   *
```

### S1PR2-LgBiT in the construct pTRE3G-BI/S1PR2-LgBiT:SmBiT-ARRB2

```
   1 ATG GGC AGC TTG TAC TCG GAG TAC CTG AAC CCC AAC AAG GTC CAG GAA CAC TAT AAT TAT ACC AAG GAG ACG CTG
     TAC CCG TCG AAC ATG AGC CTC ATG GAC TTG GGG TTG TTC CAG GTC CTT GTG ATA TTA ATA TGG TTC CTC TGC GAC
      M   G   S   L   Y   S   E   Y   L   N   P   N   K   V   Q   E   H   Y   N   Y   T   K   E   T   L

  76 GAA ACG CAG GAG ACG ACC TCC CGC CAG GTG GCC TCG GCC TTC ATC GTC ATC CTC TGT TGC GCC ATT GTG GTG GAA
     CTT TGC GTC CTC TGC TGG AGG GCG GTC CAC CGG AGC CGG AAG TAG CAG TAG GAG ACA ACG CGG TAA CAC CAC CTT
      E   T   Q   E   T   T   S   R   Q   V   A   S   A   F   I   V   I   L   C   C   A   I   V   V   E

 151 AAC CTT CTG GTG CTC ATT GCG GTG GCC CGA AAC AGC AAG TTC CAC TCG GCA ATG TAC CTG TTT CTG GGC AAC CTG
     TTG GAA GAC CAC GAG TAA CGC CAC CGG GCT TTG TCG TTC AAG GTG AGC CGT TAC ATG GAC AAA GAC CCG TTG GAC
      N   L   L   V   L   I   A   V   A   R   N   S   K   F   H   S   A   M   Y   L   F   L   G   N   L

 226 GCC GCC TCC GAT CTA CTG GCA GGC GTG GCC TTC GTA GCC AAT ACC TTG CTC TCT GGC TCT GTC ACG CTG AGG CTG
     CGG CGG AGG CTA GAT GAC CGT CCG CAC CGG AAG CAT CGG TTA TGG AAC GAG AGA CCG AGA CAG TGC GAC TCC GAC
      A   A   S   D   L   L   A   G   V   A   F   V   A   N   T   L   L   S   G   S   V   T   L   R   L

 301 ACG CCT GTG CAG TGG TTT GCC CGG GAG GGC TCT GCC TTC ATC ACG CTC TCG GCC TCT GTC TTC AGC CTC CTG GCC
     TGC GGA CAC GTC ACC AAA CGG GCC CTC CCG AGA CGG AAG TAG TGC GAG AGC CGG AGA CAG AAG TCG GAG GAC CGG
      T   P   V   Q   W   F   A   R   E   G   S   A   F   I   T   L   S   A   S   V   F   S   L   L   A

 376 ATC GCC ATT GAG CGC CAC GTG GCC ATT GCC AAG GTC AAG CTG TAT GGC AGC GAC AAG AGC TGC CGC ATG CTT CTG
     TAG CGG TAA CTC GCG GTG CAC CGG TAA CGG TTC CAG TTC GAC ATA CCG TCG CTG TTC TCG ACG GCG TAC GAA GAC
      I   A   I   E   R   H   V   A   I   A   K   V   K   L   Y   G   S   D   K   S   C   R   M   L   L

 451 CTC ATC GGG GCC TCG TGG CTC ATC TCG CTG GTC CTC GGT GGC CTG CCC ATC CTT GGC TGG AAC TGC CTG GGC CAC
     GAG TAG CCC CGG AGC ACC GAG TAG AGC GAC CAG GAG CCA CCG GAC GGG TAG GAA CCG ACC TTG ACG GAC CCG GTG
      L   I   G   A   S   W   L   I   S   L   V   L   G   G   L   P   I   L   G   W   N   C   L   G   H

 526 CTC GAG GCC TGC TCC ACT GTC CTG CCT CTC TAC GCC AAG CAT TAT GTG CTG TGC GTG GTG ACC ATC TTC TCC ATC
     GAG CTC CGG ACG AGG TGA CAG GAC GGA GAG ATG CGG TTC GTA ATA CAC GAC ACG CAC CAC TGG TAG AAG AGG TAG
      L   E   A   C   S   T   V   L   P   L   Y   A   K   H   Y   V   L   C   V   V   T   I   F   S   I

 601 ATC CTG TTG GCC ATC GTG GCC CTG TAC GTG CGC ATC TAC TGC GTG GTC CGC TCA AGC CAC GCT GAC ATG GCC GCC
     TAG GAC AAC CGG TAG CAC CGG GAC ATG CAC GCG TAG ATG ACG CAC CAG GCG AGT TCG GTG CGA CTG TAC CGG CGG
      I   L   L   A   I   V   A   L   Y   V   R   I   Y   C   V   V   R   S   S   H   A   D   M   A   A

 676 CCG CAG ACG CTA GCC CTG CTC AAG ACG GTC ACC ATC GTG CTA GGC GTC TTT ATC GTC TGC TGG CTG CCC GCC TTC
     GGC GTC TGC GAT CGG GAC GAG TTC TGC CAG TGG TAG CAC GAT CCG CAG AAA TAG CAG ACG ACC GAC GGG CGG AAG
      P   Q   T   L   A   L   L   K   T   V   T   I   V   L   G   V   F   I   V   C   W   L   P   A   F

 751 AGC ATC CTC CTT CTG GAC TAT GCC TGT CCC GTC CAC TCC TGC CCG ATC CTC TAC AAA GCC CAC TAC TTT TTC GCC
     TCG TAG GAG GAA GAC CTG ATA CGG ACA GGG CAG GTG AGG ACG GGC TAG GAG ATG TTT CGG GTG ATG AAA AAG CGG
      S   I   L   L   L   D   Y   A   C   P   V   H   S   C   P   I   L   Y   K   A   H   Y   F   F   A

 826 GTC TCC ACC CTG AAT TCC CTG CTC AAC CCC GTC ATC TAC ACG TGG CGC AGC CGG GAC CTG CGG CGG GAG GTG CTT
     CAG AGG TGG GAC TTA AGG GAC GAG TTG GGG CAG TAG ATG TGC ACC GCG TCG GCC CTG GAC GCC GCC CTC CAC GAA
      V   S   T   L   N   S   L   L   N   P   V   I   Y   T   W   R   S   R   D   L   R   R   E   V   L

 901 CGG CCG CTG CAG TGC TGG CGG CCG GGG GTG GGG GTG CAA GGA CGG AGG CGG GGC GGG ACC CCG GGC CAC CAC CTC
     GCC GGC GAC GTC ACG ACC GCC GGC CCC CAC CCC CAC GTT CCT GCC TCC GCC CCG CCC TGG GGC CCG GTG GTG GAG
      R   P   L   Q   C   W   R   P   G   V   G   V   Q   G   R   R   R   G   G   T   P   G   H   H   L

 976 CTG CCA CTC CGC AGC TCC AGC TCC CTG GAG AGG GGC ATG CAC ATG CCC ACG TCA CCC ACG TTT CTG GAG GGC AAC
     GAC GGT GAG GCG TCG AGG TCG AGG GAC CTC TCC CCG TAC GTG TAC GGG TGC AGT GGG TGC AAA GAC CTC CCG TTG
      L   P   L   R   S   S   S   S   L   E   R   G   M   H   M   P   T   S   P   T   F   L   E   G   N

                 AgeI
1051 ACG GTG GTC ACC GGT ACC GGC GGA GGG TCT AGC AGT GGC GGT GGG ATG GTC TTC ACA CTC GAA GAT TTC GTT GGG
     TGC CAC CAG TGG CCA TGG CCG CCT CCC AGA TCG TCA CCG CCA CCC TAC CAG AAG TGT GAG CTT CTA AAG CAA CCC
      T   V   V   T   G   T   G   G   G   S   S   S   G   G   G   M   V   F   T   L   E   D   F   V   G

1126 GAC TGG GAA CAG ACA GCC GCC TAC AAC CTG GAC CAA GTC CTT GAA CAG GGA GGT GTG TCC AGT TTG CTG CAG AAT
     CTG ACC CTT GTC TGT CGG CGG ATG TTG GAC CTG GTT CAG GAA CTT GTC CCT CCA CAC AGG TCA AAC GAC GTC TTA
      D   W   E   Q   T   A   A   Y   N   L   D   Q   V   L   E   Q   G   G   V   S   S   L   L   Q   N

1201 CTC GCC GTG TCC GTA ACT CCG ATC CAA AGG ATT GTC CGG AGC GGT GAA AAT GCC CTG AAG ATC GAC ATC CAT GTC
     GAG CGG CAC AGG CAT TGA GGC TAG GTT TCC TAA CAG GCC TCG CCA CTT TTA CGG GAC TTC TAG CTG TAG GTA CAG
```

L A V S V T P I Q R I V R S G E N A L K I D H V

1276 ATC ATC CCG TAT GAA GGT CTG AGC GCC GAC CAA ATG GCC CAG ATC GAA GAG GTG TTT AAG GTG GTG TAC CCT GTG

TAG TAG GGC ATA CTT CCA GAC TCG CGG CTG GTT TAC CGG GTC TAG CTT CTC CAC AAA TTC CAC CAC ATG GGA CAC

I I P Y E G L S A D Q M A Q I E E V F K V V Y P V

1351 GAT GAT CAT CAC TTT AAG GTG ATC CTG CCC TAT GGC ACA CTG GTA ATC GAC GGG GTT ACG CCG AAC ATG CTG AAC

CTA CTA GTA GTG AAA TTC CAC TAG GAC GGG ATA CCG TGT GAC CAT TAG CTG CCC CAA TGC GGC TTG TAC GAC TTG

D D H H F K V I L P Y G T L V I D G V T P N M L N

1426 TAT TTC GGA CGG CCG TAT GAA GGC ATC GCC GTG TTC GAC GGC AAA AAG ATC ACT GTA ACA GGG ACC CTG TGG AAC

ATA AAG CCT GCC GGC ATA CTT CCG TAG CGG CAC AAG CTG CCG TTT TTC TAG TGA CAT TGT CCC TGG GAC ACC TTG

Y F G R P Y E G I A V F D G K K I T V T G T L W N

1501 GGC AAC AAA ATT ATC GAC GAG CGC CTG ATC ACC CCC GAC GGC TCC ATG CTG TTC CGA GTA ACC ATC AAC AGT TAA

CCG TTG TTT TAA TAG CTG CTC GCG GAC TAG TGG GGG CTG CCG AGG TAC GAC AAG GCT CAT TGG TAG TTG TCA ATT

G N K I I D E R L I T P D G S M L F R V T I N S *

**Fig. S2.** The nucleotide and amino acid sequences of the untagged or the C-terminally LgBiT-tagged human S1PR2. The amino acid sequence of human S1PR2 is shown in red and that of LgBiT in blue. A synonymous nucleotide mutation of human S1PR2 compared to the reference sequence (NM_004230) is shown in red and shaded. The restriction enzyme cleavage sites for gene cloning are shaded.

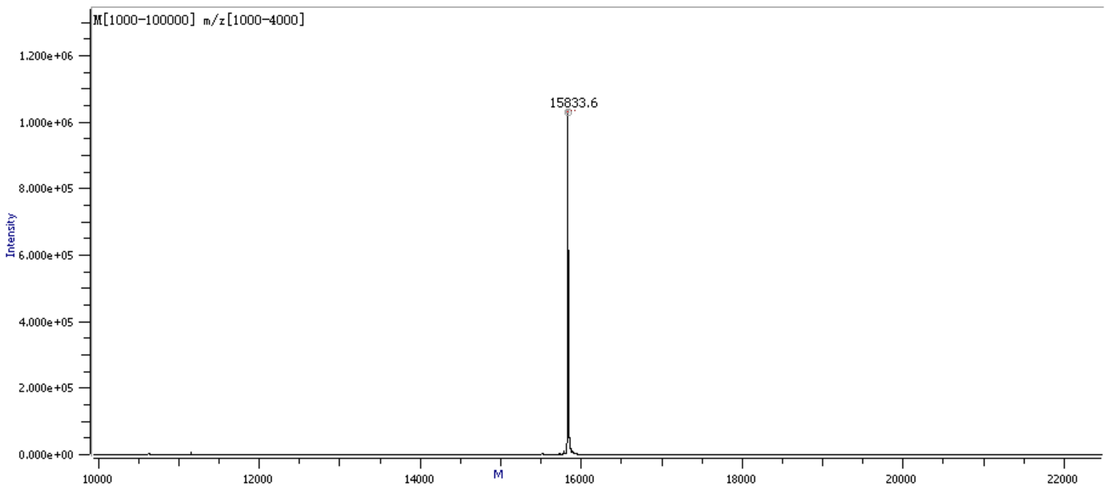


**Fig. S3.** Molecular mass of the recombinant mature human MYDGF measured on an AB SCIEX TripleTOF 5600 mass spectrometer. Its theoretical molecular mass is 15832.6.

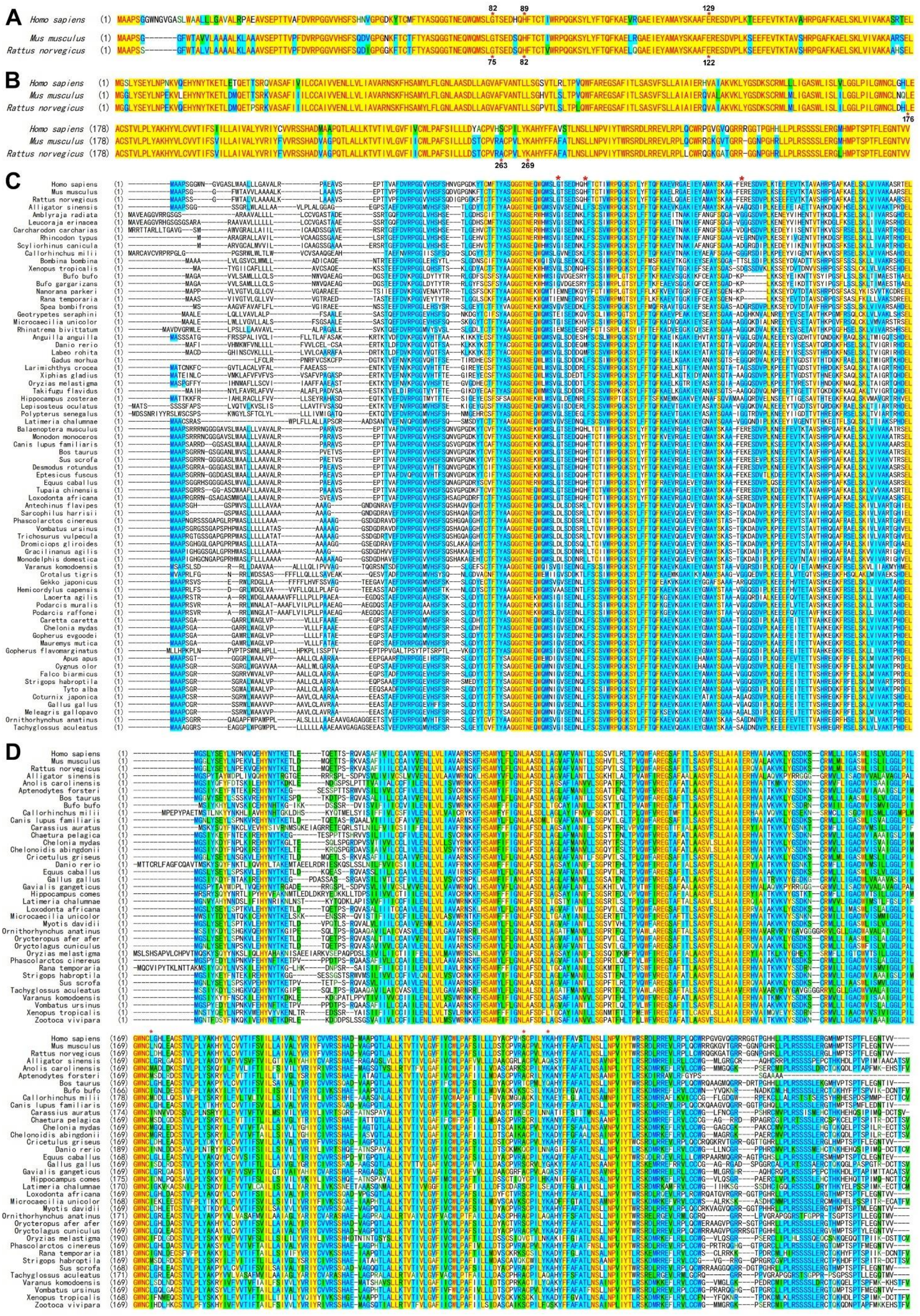
A
B
C
D

**Fig. S4.** Amino acid sequence alignment of MYDGF orthologs and S1PR2 orthologs from different vertebrate species. (**A**) Amino acid sequence alignment of MYDGF orthologs from human, mouse, and rat. (**B**) Amino acid sequence alignment of S1PR2 orthologs from human, mouse, and rat. (**C**) The amino acid sequence alignment of MYDGF orthologs from mammals, birds, reptiles, amphibians, and fishes. (**D**) The amino acid sequence alignment of S1PR2 orthologs from mammals, birds, reptiles, amphibians, and fishes. Positions of the so-called receptor-binding residues in MYDGF orthologs and ligand-binding residues in S1PR2 orthologs proposed by the recent paper [13] are indicated by red asterisks.
